# The cold tolerance of the terrestrial slug, *Ambigolimax valentianus*

**DOI:** 10.1101/2024.10.29.620936

**Authors:** Lauren T. Gill, Hiroko Udaka, Katie E. Marshall

## Abstract

Terrestrial molluscs living in temperate and polar environments must contend with cold winter temperatures. However, the physiological mechanisms underlying the survival of terrestrial molluscs in cold environments and the strategies employed by them are poorly understood. Here we investigated the cold tolerance of *Ambigolimax valentianus*, an invasive, terrestrial slug that has established populations in Japan, Canada, and Europe. To do this, we acclimated *A. valentianus* to different environmental conditions (differing day lengths and temperatures), then exposed them to sub-zero temperatures and measured overall survival. Then, we measured low molecular weight metabolites using ^1^H NMR to see if they play a role in their cold tolerance as they do in other invertebrate species. We found that *A. valentianus* is not strongly freeze tolerant but do become more cold-hardy after acclimation to shorter day lengths. We also found that no metabolites were strongly upregulated in response to winter conditions despite the change in cold hardiness, and instead saw evidence of metabolic suppression leading up to winter such as formate and L-glutamine being supressed in winter conditions.

## Introduction

Invertebrates living on land must effectively navigate the daily and seasonal variations of terrestrial environments. Among these, terrestrial molluscs inhabiting temperate and polar regions must also have a survival mechanism to tolerate winter temperatures while also maintaining a moist body surface for respiration (Prior et al., 1983). Terrestrial slugs species may be particularly susceptible to low winter temperatures as they do not have shells or epiphragms like those of land snails, so the moist skin on their body surface can come in direct contact with ice crystals on the ground, initiating inoculative freezing of their body fluids (Ansart & Vernon, 2003; Udaka et al., 2008). Despite the growing literature on invertebrate cold and freeze tolerance, many questions about the abilities of terrestrial molluscs to tolerate low temperatures and freezing remain unanswered.

Clear distinctions between freeze tolerance and other forms of cold tolerance are critical for understanding the winter survival strategies of gastropods, but these are often not distinguished in the literature (Gill et al., 2024). Cold tolerance is the ability to survive and function at low temperatures whereas freeze tolerance is the ability to survive the freezing of body fluids. Importantly, an organism that is cold tolerant is not necessarily freeze tolerant, and vice versa. For example, the cold tolerant spruce budworm *Choristoneura fumiferana* can survive temperatures of -35 °C or lower but will die upon freezing (Butterson et al., 2021). In contrast, the freeze tolerant bay mussel *Mytilus trossulus* can survive freezing for several hours but dies when temperatures drop below -10 °C (Kennedy et al., 2020). Given that freeze tolerance is a distinct survival mechanism, distinguishing it from general cold tolerance is crucial for understanding how gastropods tolerate winter temperatures. Measuring an organism’s body temperature during a cold exposure can reveal whether freezing has occurred (through observing a supercooling point and subsequent heat release) providing more insight into the strategy an animal uses when faced with sub-zero temperatures (Sinclair et al., 2015).

In response to cold winter temperatures, terrestrial molluscs typically employ one of two strategies to survive: partial tolerance to freezing or behavioural changes such as moving to a new habitat. Partial freeze tolerance refers to an animal’s ability to survive when a portion of their body water is converted to ice (Sinclair, 1999). Because ice formation is an exothermic process, an animal’s body will be warmer than the environment when ice initially forms due to the heat production by the latent heat of crystallization. Animals that exhibit partial freeze tolerance will only survive if their body is warmed before their body temperature equilibrates to the environmental temperature (Sinclair, 1999). For example, the partially freeze tolerant slug *Deroceras reticulatum* can tolerate freezing for up to 30 minutes before facing mortality (R. Cook, 2004). Whether terrestrial slugs display freeze tolerance appears to be species-specific. There are several slug species that do not survive freezing, including *Limacus flavas, D. altaicum,* and *Arion circumscriptus* (Berman et al., 2011; Riddle, 1983). However, *D. reticulatum*, *D. laeve,* and slugs of the genus *Arion* can tolerate freezing (R. Cook, 2004; Slotsbo et al., 2012).

Some terrestrial molluscs may also exhibit behavioural changes when faced with sub-zero temperatures, for example by secreting a layer of mucus to seal themselves into their shell, evacuating their gut contents, or moving to a warmer site that buffers cold temperatures to prevent inoculative freezing (Ansart & Vernon, 2003; Douglas & Tooker, 2012). Adult slugs will often overwinter in a thermally buffered site, for example under crop residue or leaf litter (Douglas & Tooker, 2012). Slugs have also been known to display ‘huddling’ behaviour in response to cold temperatures, likely to prevent evaporative water loss from their body surface (A. Cook, 1981; R. Cook, 2004; Riddle, 1983). All of these strategies either keep the organism’s body fluids above the freezing point during the winter or prevent external and internal ice nucleation. Terrestrial molluscs that are not freeze tolerant can adopt these behavioural strategies to avoid freezing, enabling their survival during the cold winter months.

Many terrestrial molluscs exhibit a seasonality in their temperature tolerances, becoming more heat tolerant in the summer months, and more cold and freeze tolerant in the winter months (Udaka et al., 2008). Cooling temperatures and shortening days are two environmental factors that can signal that winter temperatures are approaching to a terrestrial mollusc (Ansart & Vernon, 2003). Low temperature acclimation has been shown to significantly affect cold hardiness in several species of snails and slugs (Aarset, 1982; R. Cook, 2004; Udaka et al., 2008). For example, *D. reticulatum* slugs collected in the winter survived better during freeze exposures than those collected in the summer (R. Cook, 2004). Similarly, after low temperature acclimation, lab-reared *Ambigolimax valentianus* slugs were better able to survive short cold exposures (Udaka et al., 2008). Photoperiod also affects cold and freeze tolerance. In the summer, days are typically longer and, in the winter, days are typically shorter. Seasonal day length differences are most pronounced near the poles, and this region is also where mollusc species are likely to experience the coldest winters. Decreasing day length has been shown to increase cold tolerance of both slug and snail species (Aarset, 1982; Ansart & Vernon, 2003; Udaka et al., 2008). Finally, the body size of a mollusc affects its cold tolerance. The brown garden snail, *Cornu aspersu,* and another large species of snail, *Arianta arborustorum*, are both partially freeze tolerant (Ansart & Vernon, 2003). In contrast, small snails like *Anguispira alternata*, *Discus cronkhitei*, *Gastrocopta armifera* and *Vallonia perspectiva* are all freeze avoidant (Ansart & Vernon, 2003). Larger organisms are at a greater risk of freezing, as the probability of ice formation increases with an increasing number of water molecules (Bigg, 1953; Zachariassen & Kristiansen, 2000). This explains why larger species may display freeze tolerance more often and suggests the body size of a species (small vs. large) can determine which strategy an animal employs in response to freezing conditions (Ansart & Vernon, 2003).

Terrestrial molluscs exhibit an array of metabolic responses to cold temperatures. Some freeze tolerant slugs generate metabolites like lactate and succinate to anaerobically support O_2_-depleted tissues during freezing (Slotsbo et al., 2012). Another important metabolite is glutamine, an amino acid and metabolic intermediate that has many important functions in the body (Labow & Souba, 2000). In *D. reticulatum*, a slug species introduced to North America from Europe, glutamine levels strongly decrease after freezing, while glucose and aspartate levels remained unchanged (Storey et al., 2007). The native counterpart to *D. reticulatum*, *D. laeve,* also downregulates glutamine in response to freezing (Storey et al., 2007). This suggests that molluscs may alter their metabolite profiles, including small molecules like amino acids, in response to cold temperatures.

Many freeze-tolerant and cold-tolerant organisms accumulate molecules called cryoprotectants to help protect cells and tissues in sub-zero temperatures (Lee, 2010; Toxopeus & Sinclair, 2018). Cryoprotectants can be categorized into two primary groups based on their molecular weight: high molecular weight cryoprotectants which include antifreeze proteins and ice-nucleating agents, and low molecular weight cryoprotectants (Duman, 2001; Zachariassen, 1992). Low molecular weight cryoprotectants are small metabolites such as amino acids and sugars that can have a variety of cryoprotective functions. Low molecular weight cryoprotectants can decrease the ice content in the tissue by depressing the freezing point and can also directly protect membranes and proteins (Storey & Storey, 1996). Animals can upregulate these protective mechanisms before and during freezing or repair the cellular damage after the freeze exposure. Insect species are able to accumulate low molecular weight cryoprotectants in high concentrations to avoid freezing damage (Lee, 2010). In intertidal molluscs, accumulating relatively small quantities of low molecular weight cryoprotectants is associated with greater freeze tolerance (Kennedy et al., 2020). Terrestrial molluscs can also change concentrations of low molecular weight cryoprotectants in response to cold or freezing stress. Scandinavian slugs from the genus *Arion* upregulate glucose, lactate, and succinate after freezing (Slotsbo et al., 2012). Low molecular weight cryoprotectant accumulation appears to be species-specific but may play a role in freeze tolerance in some slug species.

The slug *Ambigolimax valentianus* is indigenous to the Iberian peninsula in Europe but has since been introduced widely throughout many regions of the world including North America, South America, the United Kingdom, and Japan (Satoh et al., 2020; Walden, 1962). This invasive species of slug inhabits greenhouses and anthropogenically-altered areas and lowers the value of horticulture and agriculture products by feeding and leaving mucus trails, similar to many other invasive terrestrial molluscs (Cowie et al., 2009; Satoh et al., 2020). Slugs of the species *A. valentianus* eat plant and animal matter (Kurozumi, 2002; unpublished observations). They are a monoecious species; one individual has both male and female reproductive organs, with mature sperm produced earlier in the year from September – April, and oocytes produced later from October – May (Udaka et al., 2007). *Ambigolimax valentianus* lay their eggs from November – May, but hatching success is lowered in the coldest winter months (January and February in Japan; Udaka et al., 2007).

Slug species like *A. valentianus* are at risk of freezing during the winter if they come in direct contact with ice crystals on the ground (Udaka et al., 2008). *Ambigolimax valentianus* can survive sub-zero temperatures and can modulate their cold tolerance seasonally (Udaka et al., 2008). For example, *A. valentianus* reared under long day (LD; 16L:8D) conditions at 20 °C had a 25% survival rate when exposed to a short term cold exposure at -7 °C (Udaka et al., 2008). By contrast, slugs reared under short day (SD; 12L:12D) conditions at 15 °C, almost all survive the same -7 °C cold exposure (Udaka et al., 2008). However, it remains unclear whether the slugs in this study froze, as the supercooling point—which would indicate whether freezing occurred—was not measured. The supercooling point (SCP) is defined as the lowest temperature an organism reaches before the onset of an exothermic reaction, where body fluids begin to freeze. In other terrestrial mollusc species, supercooling points have been recorded within a range of -1 °C to -24 °C (Ansart & Vernon, 2003). While it is established that *A. valentianus* can tolerate sub-zero temperatures of up to -7 °C, whether it can tolerate freezing is unknown.

There are many unknowns surrounding the physiological and biochemical impacts of freezing and cold exposure in the invasive species *A. valentianus*: the degree to which *A. valentianus* has the ability to survive internal ice formation, the unknown effects of environmental factors on its cold hardiness, and finally the poorly-understood role of high or low cryoprotectants in the cold tolerance of *A. valentianus*. It’s important to investigate these questions as invasive species often exhibit broader thermal tolerances, and likely enhanced lower thermal tolerance thresholds compared to their native counterparts (Kelley, 2014) enabling them to survive in a wider range of temperatures and facilitating their spread into new environments. Consequently, the dispersal patterns of invasive molluscs will often be set by abiotic factors such as temperature, and thus understanding the temperature tolerances of these species can help to prevent further establishment in new regions (Satoh & Yamazaki, 2020).

The first objective of this study is to understand how environmental factors like photoperiod (day length) and cold can result in changes in survival and supercooling point (the temperature at which ice begins to form in the body). To do this, we acclimated slugs to fully factorial combinations of two different temperatures (15 °C or 20 °C) and two different photoperiods (short day or long day), measuring their supercooling points and survival, with the expectation that colder temperatures and shorter photoperiods would enhance low temperature survival and alter supercooling points. Also, by exposing *A. valentianus* to sub-zero temperatures and subsequently evaluating survival, we aimed to determine its freeze tolerance, predicting it would be partially freeze tolerant similar to other terrestrial slugs (e.g., *D. reticulatum*; Cook, 2004). The second objective of this study is to examine how photoperiod and temperature change metabolite concentrations and how those changes relate to low temperature survival. We used a subset of acclimated slugs to conduct NMR spectroscopy to quantify metabolites and putative low molecular weight cryoprotectants, predicting that winter acclimation would lead to low molecular weight cryoprotectant accumulation.

## Methods

### *Ambigolimax valentianus* collection and acclimation

Specimens of *Ambigolimax valentianus* were collected from Kyoto University Campus, Kyoto Prefecture, Japan (35.02928° N, 135.78285° W) between June 21 and July 10, 2023. Animals collected were selected for an approximate body weight of 0.5-1.0 grams. Within 30 minutes of collection, slugs were placed in plastic containers lined with moist paper towel (Fig. S1) in incubators (Panasonic MIR-154-PJ or MIR-154S-PJ) set to long day conditions (16:8 L:D) at 20 °C. Slugs were fed insect food and fresh carrots. After seven days of laboratory acclimation, the slugs were divided and placed in one of four conditions: long day at 20 °C, short day at 20 °C, long day at 15 °C, or short day at 15°C. At least 15 slugs were placed in each condition, to achieve a sample size of 12 for the following experiment. These day lengths and temperatures were based on Udaka et al. (2018), which used two of the same acclimation conditions and found that *A. valentianus* had significantly enhanced cold tolerance at short day 15°C conditions. Slugs acclimated at long day conditions were exposed to 16h light and 8h darkness, and slugs at short day conditions were exposed to 12h light and 12h darkness. These experimental photoperiods reflect Kyoto, Japan’s summer and winter day lengths (∼15.5 hours of light on the longest day of the year and ∼11 hours of daylight on the shortest day of the year; https://www.timeanddate.com/sun/japan/kyoto?). Slugs were kept in these new conditions for 21 days before sampling.

## Freezing

### Testing effects of acclimation on freezing tolerance

Each slug was weighed using an analytical balance (AB54-S, Mettler Toledo; with a sensitivity of 0.001 g) and then was placed in a 1.5 mL microcentrifuge tube with a small hole in the bottom to allow for air circulation. One or two type-T thermocouples were then placed in contact with the body of the slug and the tube was plugged with cotton to keep the slug and thermocouples in place. The thermocouples were then taped in place using electrical tape (Fig. S2). Thermocouples were connected to computers equipped with Graphtec GL100_240_840_APS Version 1.30 for Windows through a Graphtec GL100 interface throughout the freeze exposure to track the slug’s body temperatures and record freezing events. Freezing events were detected as a rapid release of heat and dramatic rise in body temperature, which happens immediately after the supercooling point is reached (which is the point where body fluids begin to turn to ice; Lee, 2010). To initiate the assay, the tube containing the slugs then placed in a two double-lined plastic bags which were suspended in a programmable cooling bath (LTB-250A Low Temperature Water Bath, As One) with an initial temperature of 5 °C, then cooled at a rate of 0.75°C/minute. Once the SCP had been observed and temperature plateaued (indicated by red arrow in Fig. S3), the tube containing the slug was immediately removed from the cooling bath, and the cotton and thermocouples were removed. The tubes and slugs were then immediately placed in a Petri dish lined with moist paper towel in a 15 °C incubator and allowed to recover for 24 hours before survival was determined.

### Comparing survival of cold exposed and freeze exposed slugs

To clearly distinguish between the effects of cold exposure and freezing exposure, a subset of 11 slugs that had been acclimated to long-day, 15°C conditions were selected. Slugs from this particular acclimation condition were chosen purely for logistical reasons: the experiment was planned after acclimation had already begun, and we had a surplus of slugs acclimated to this condition. These 11 slugs were exposed to -6 °C (the average temperature of their supercooling point as determined in prior trials) for around 35 minutes. If a SCP occurred, it was recorded. The tubes and slugs were then immediately placed in a Petri dish lined with moist paper towel in a 15 °C incubator and allowed to recover for 24 hours before survival was determined. The exact duration of time each slug spent in the freezing bath varied slightly from 35 minutes, as they were removed one at a time, but the specific time for each slug was recorded.

### Metabolite assays

Slugs for ^1^H NMR were collected between July 19 and July 28, 2023. As before, slugs were acclimated for one week at LD 20 and then three weeks at either long day at 20 °C, short day at 20 °C, long day at 15 °C, or short day at 15 °C. After acclimation, slugs were weighed, placed in a 1.5 mL microcentrifuge tube, and snap frozen with liquid nitrogen. The slugs were then lyophilized using a Taitec VD-250F Freeze dryer for 24 hours, transported back to Vancouver in freezer boxes (Heathrow Scientific True North Mini-Cooler) over the course of 48 hours, and finally stored at -20 °C until further use. 1-D proton nuclear magnetic resonance spectroscopy (NMR) was carried out using methods similar to (Kennedy et al., 2020; Slotsbo et al., 2012). The freeze-dried slugs were weighed to record their dry mass then crushed into a powder using a bullet blender (Bullet Blender 50 Blue, Next Advance) in a 4 °C fridge with 200 μL of 3.2 mm round beads for 5 min at setting 8 in 2 mL microcentrifuge vials (Eppendorf safe-lock). To extract water-soluble metabolites, 500 μL of cold 50% acetonitrile (CH_3_CN) was added to the tubes and homogenized using a bullet blender (Bullet Blender 50 Blue, Next Advance) for 10 mins at setting 8.

Homogenates were centrifuged at 7000×g for 7 min, supernatant was transferred to new 1.5 mL tubes, then dried in a centrifugal vacuum concentrator (Eppendorf 5301) and stored at −80 °C. The dried polar extracts were resuspended in 600 μl of 50 mM sodium phosphate buffer (pH 6.9, 75% deuterium oxide, Sigma-Aldrich) immediately before NMR analysis. The phosphate buffer contained 1 mM 2,2-dimethyl-2-silapentane-5-sulfonate (DSS; Sigma-Aldrich) as internal reference. The samples were vortexed and then 500 μL was transferred to a 5 mm NMR tube for ^1^H NMR.

A Bruker Advance spectrometer was used to perform ^1^H NMR data acquisition. One-dimensional ^1^HNMR spectra were acquired at a frequency of 600.15 MHz at an 8.4 kHz spectral width, with 64 scans at 300 K, using TopSpin software (v 2.1, Bruker) with water suppression (a relaxation delay of 3 seconds and a power of 50dB that corresponds to a field strength of around 25Hz) requiring 10 min of acquisition time. Determination of metabolite concentrations was performed using Bayesil (Ravanbakhsh et al., 2015), software that conducts fully automated spectral processing and profiling. Metabolite profiling was completed using the Human Metabolome Database compound spectral reference library (Wishart et al., 2018) and selecting the serum biofluid and the standard profiling speed. The Bayesil pipeline consists of initial spectral processing, spectral deconvolution using algorithms to transform sections of the spectrum into probabilistic models, then applying an approximate inference over the model to present the most probable metabolic profile (Ravanbakhsh et al., 2015). The software then determines the concentrations of individual metabolites using the concentration of the known DSS signal (1mM).

### Statistical analysis

Statistical analyses were performed using R (c. 4.3.2; R Development Core Team 2023). The R package “ggplot2” was used to generate Fig. 2, Fig. 4, Fig. 5 (Wickham, 2016), and the R package “sciplot” was used to generate Fig. 3 (Morales & Murdoch, 2020). The R package “plyr” was also used to determine means and standard errors of proportions before plotting (Wickham, 2011).

A logistic regression model was fitted to assess the relationship between the binary outcome variable (survival: alive vs. dead) and a set of predictor variables including acclimation day length (long day or short day), acclimation temperature (15 °C or 20 °C), and slug weight. The model (glm) was specified as follows: survival ∼ acclimation day length × acclimation temperature + weight, with a binomial distribution. Following the model fit, we conducted likelihood ratio tests to evaluate the significance of each predictor using the anova() function with test = “Chisq”. This approach allowed us to assess whether the inclusion of each variable significantly improved model fit. The test compares the deviance of models with and without each predictor, with the resulting p-values derived from a chi-squared distribution. Then, a Tukey post-hoc test was conducted using the “TukeyHSD” function in R to identify statistically significant differences between acclimation conditions. A similar method was used to test for significant differences in survival after cold or freeze exposures. A logistic regression model was fitted to assess the relationship between the binary outcome variable (survival: alive vs. dead) and a set of predictor variables including cold exposure length (minutes), freeze occurrence (presence or absent), and slug weight as potential predictors of survival after a cold exposure. The model was specified as follows: survival ∼ weight + cold exposure length + freeze occurrence. The resulting p-values were derived from a chi-squared distribution. Then, to test if supercooling point varied by acclimation condition, a two-way ANCOVA model was fit using the aov() function in R. The factors included in the analysis were acclimation day length (short day or long day) and acclimation temperature (15°C or 20°C), and slug weight (in grams) was included as a continuous covariate in the model to control for its potential influence on SCP. Finally, to test if body water percentage was affected by acclimation condition, a linear model was fit using acclimation day length and acclimation temperature as predictor values. Models were compared using the anova() function in R, and evaluated using an F test statistic.

To determine how metabolite content changes with different acclimation conditions, outputted metabolite concentrations (in µM) from Bayesil were transformed into molar quantities (µmol) by multiplying each quantity by the volume of diluent in sample prep (500 µL). Then, the total moles of each metabolite within a sample were normalized by slug’s wet mass. The normalized data were then square root transformed and then auto-scaled (mean-centered and divided by the standard deviation of each variable) using the R package “mdattools” (Kucheryavskiy, 2020) to achieve a normal distribution. Metabolite concentrations are reported as µmol g^−1^ wet mass.

The “Statistical Analysis [metadata table]” module on MetaboAnalyst 6.0 (https://www.metaboanalyst.ca; Xia et al., 2009) was then used to perform a two-way ANOVA on each metabolite, identifying those were statistically significant (p<0.05, corrected for False Discovery Rate). For maximum reproducibility and transparency in code, a two-way ANOVA was run again in R for those metabolites that MetaboAnalyst 6.0 identified, and outputs were recorded.

To identify any changes in metabolic pathways following different acclimation conditions, metabolite concentrations were analyzed through the “Pathway Analysis” module from MetaboAnalyst 6.0 using the *Drosophila melanogaster* pathway library (KEGG pathway info were obtained in Dec. 2023). Only data from slugs acclimated at long day 20 °C or short day 15°C were used in this analysis.

## Results

### Survival

To test the effects of acclimation on freezing tolerance of *A. valentianus*, 48 slugs were cooled until their supercooling point was reached (the temperature exotherm indicative of freezing had been observed; Fig. S3). On average, slugs took 20:36 (± 2:32) minutes to freeze. Three out of the 48 slug tubes were contaminated with the ethanol from the ethanol bath because of a leak, and thus were excluded from the survival analysis. Slugs that were acclimated to short day conditions had higher survival following this exposure than slugs that were acclimated to long day conditions (p = 0.017, df = 1, 40; Fig. 1). There was no significant effect of acclimation temperature on survival (df = 1, 40, p = 0.61) and slug mass was also not a significant predictor of survival (df = 1, 40, p = 0.12). Although not quantitatively measured, cold and short-day acclimated slugs were often observed displaying ‘huddling’ behaviour as shown in Fig. S5.

**Fig. 1:**
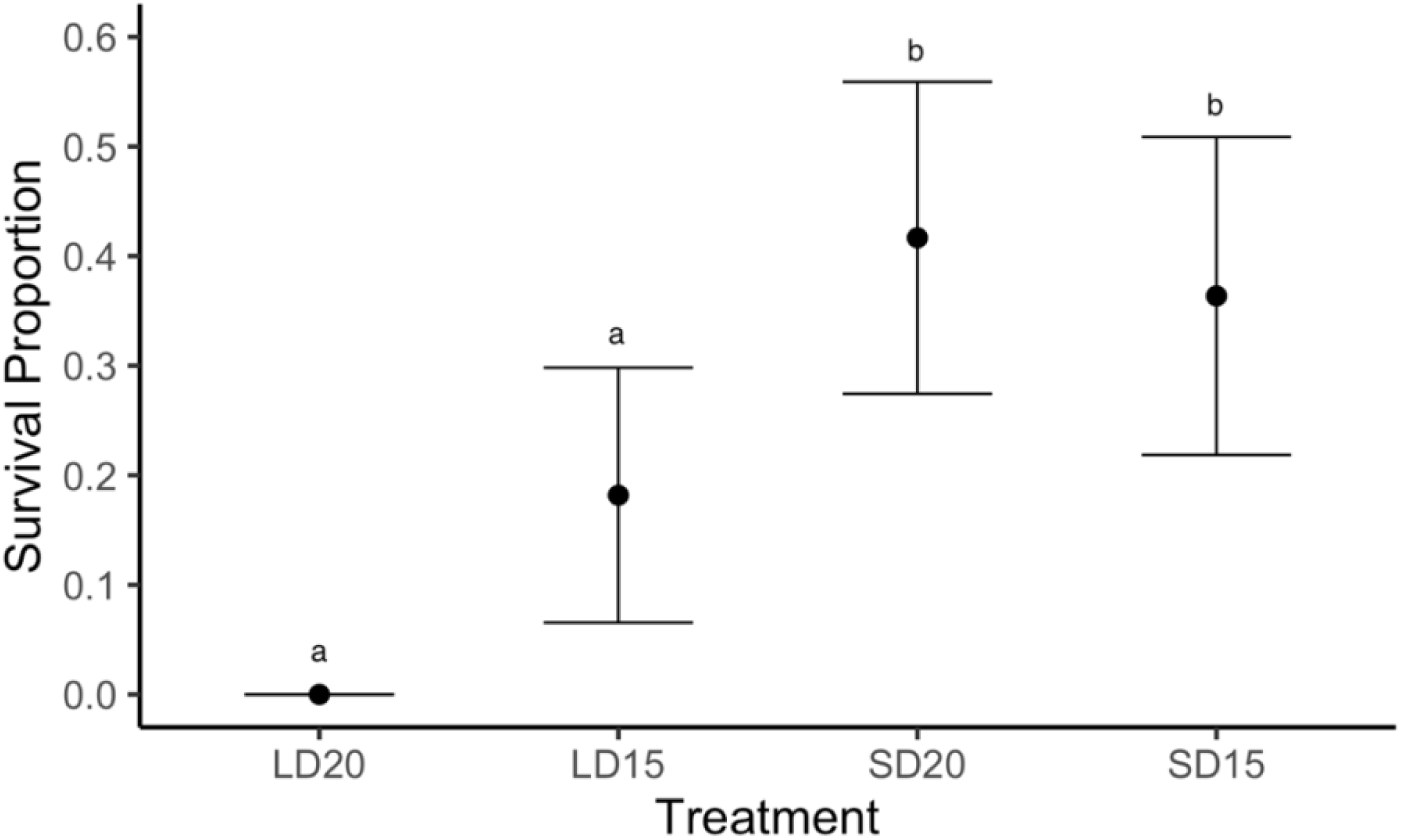
Survival proportion of *A. valentianus* immediately removed post initiation of ice (after SCP exotherm had been observed). Slugs were cooled at a rate of 0.75°C/minute. Groups with the same letter are not statistically different, error bars indicate standard error of the proportion. Treatment abbreviations are as follows: LD20 is long day 20 °C, LD15 is long day 15 °C, SD20 is short day 20 °C, and SD15 is short day 15 °C. Logistic regression p_DayLength_ = 0.017, n = 11 or 12.

To separate the effects of cold exposure versus freezing exposure on *A. valentianus*, a new subset of 11 slugs acclimated at long day 15 °C conditions were exposed to -6 °C (a temperature slightly below the average temperature of their supercooling point as determined in prior trials). Trial was conducted until approximately half of the slugs froze, which took around 35 minutes. Five of the 11 slugs froze, which was evaluated by the presence of an exotherm (as in Fig. S3) All slugs that froze during the cold exposure died, regardless of the small variations in time spent frozen (Fig. 2). All slugs that did not freeze survived.

**Fig. 2.**
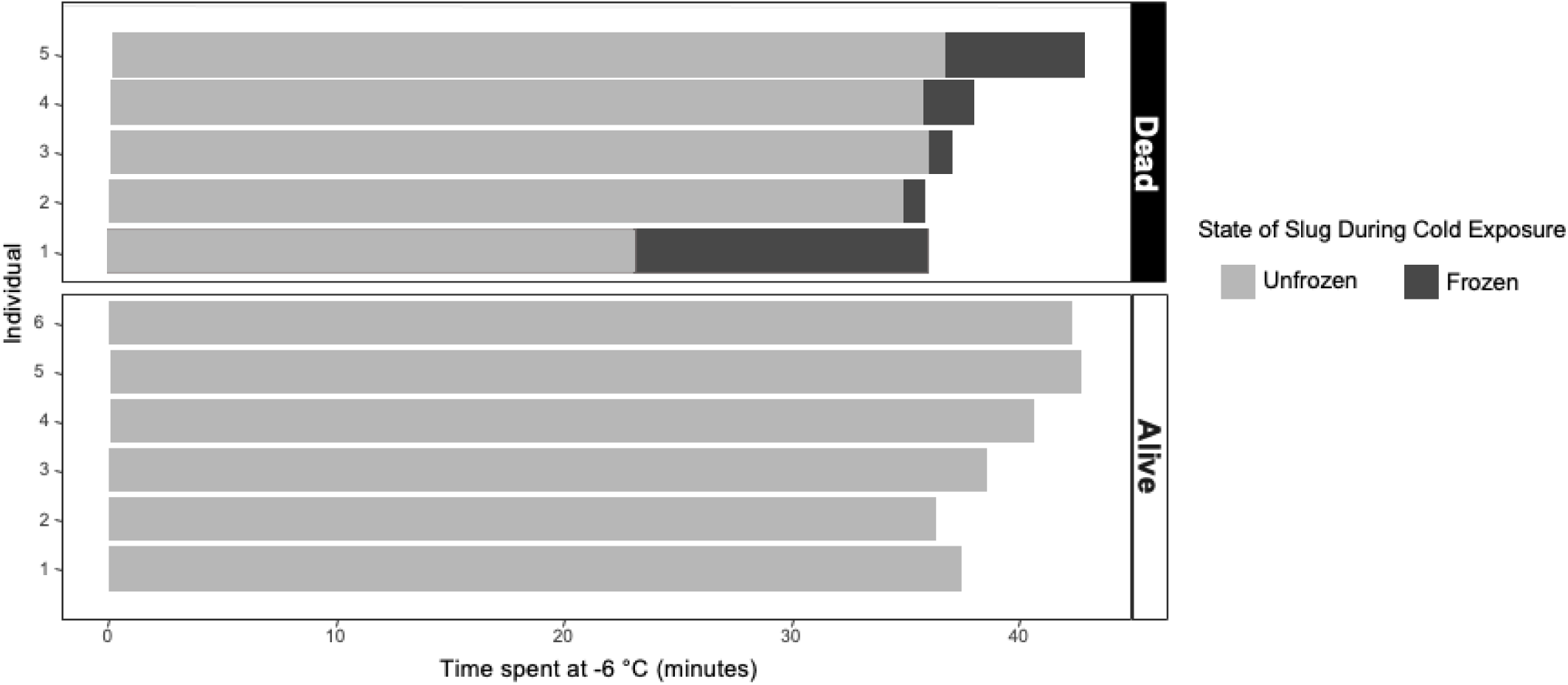
The survival of *A. valentianus* exposed to -6 °C (just below their super-cooling point) for approximately 35 minutes, and whether or not they froze during the cold exposure. Slugs were kept under long day conditions at 15 °C. N = 11, logistic regression, effect of freezing, p = 0.0002.

### Supercooling Point

There was no significant effect of acclimation temperature (df = 1, 36, p = 0.37) or acclimation day length (df = 1, 36, p = 0.32) on the supercooling point of *A. valentianus* (Fig. 3). The interaction between acclimation temperature and acclimation day length was also not significant (df = 1, 36, p = 0.59). Supercooling point temperature was affected by slug weight, with heavier slugs having a higher supercooling point (df = 1, 36, p = 0.022; Fig. 4).

**Fig. 3.**
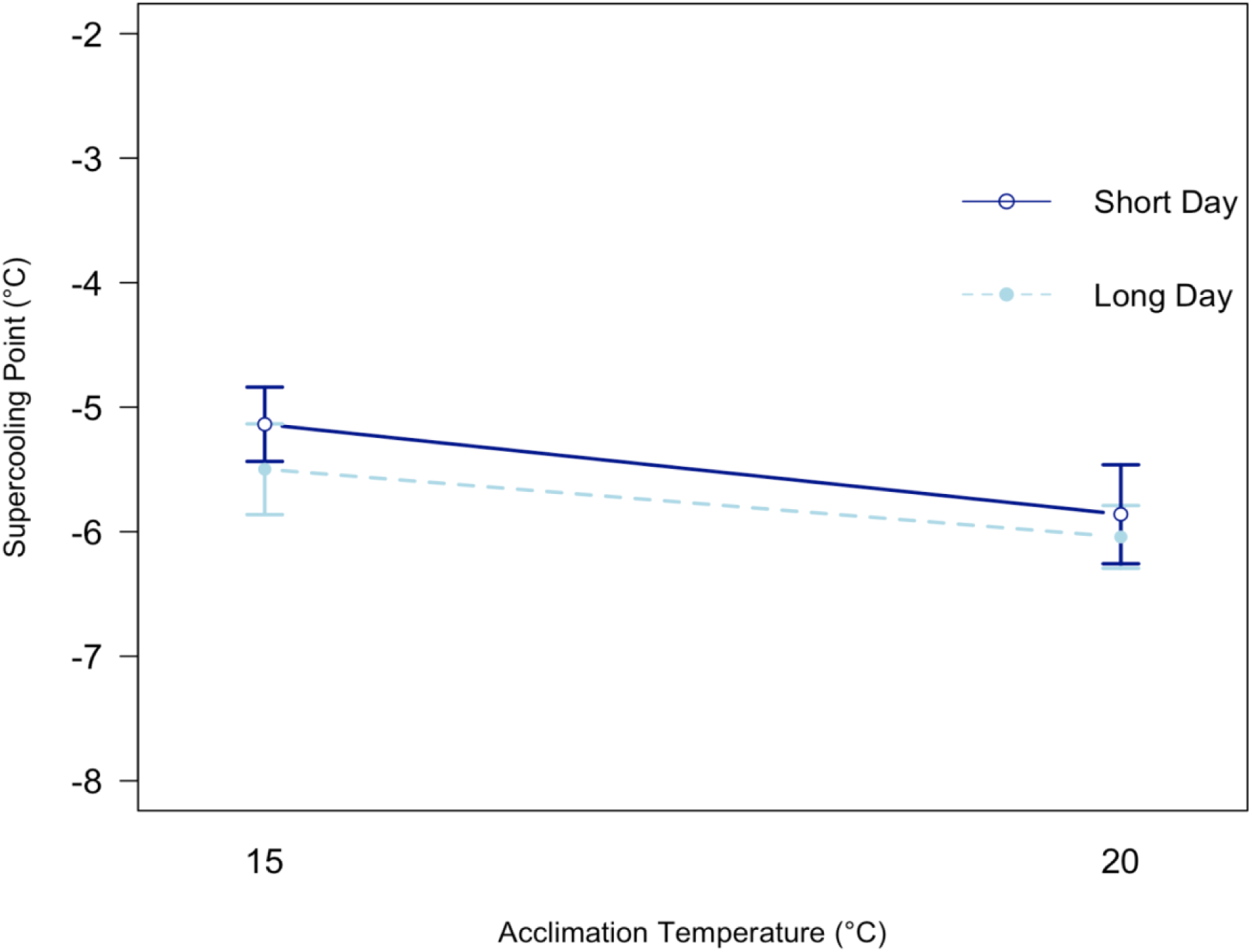
Supercooling point of *A. valentianus* does not significantly differ based on acclimation temperature or daylength. SD = Short day conditions, LD = Long day conditions. Points represent mean and error bars represent standard error. No significant difference found between treatments (acclimation temperature: p = 0.3729, acclimation day length: p = 0. 3219, temperature × day length: p = 0. 5885). n = 12.

**Fig. 4.**
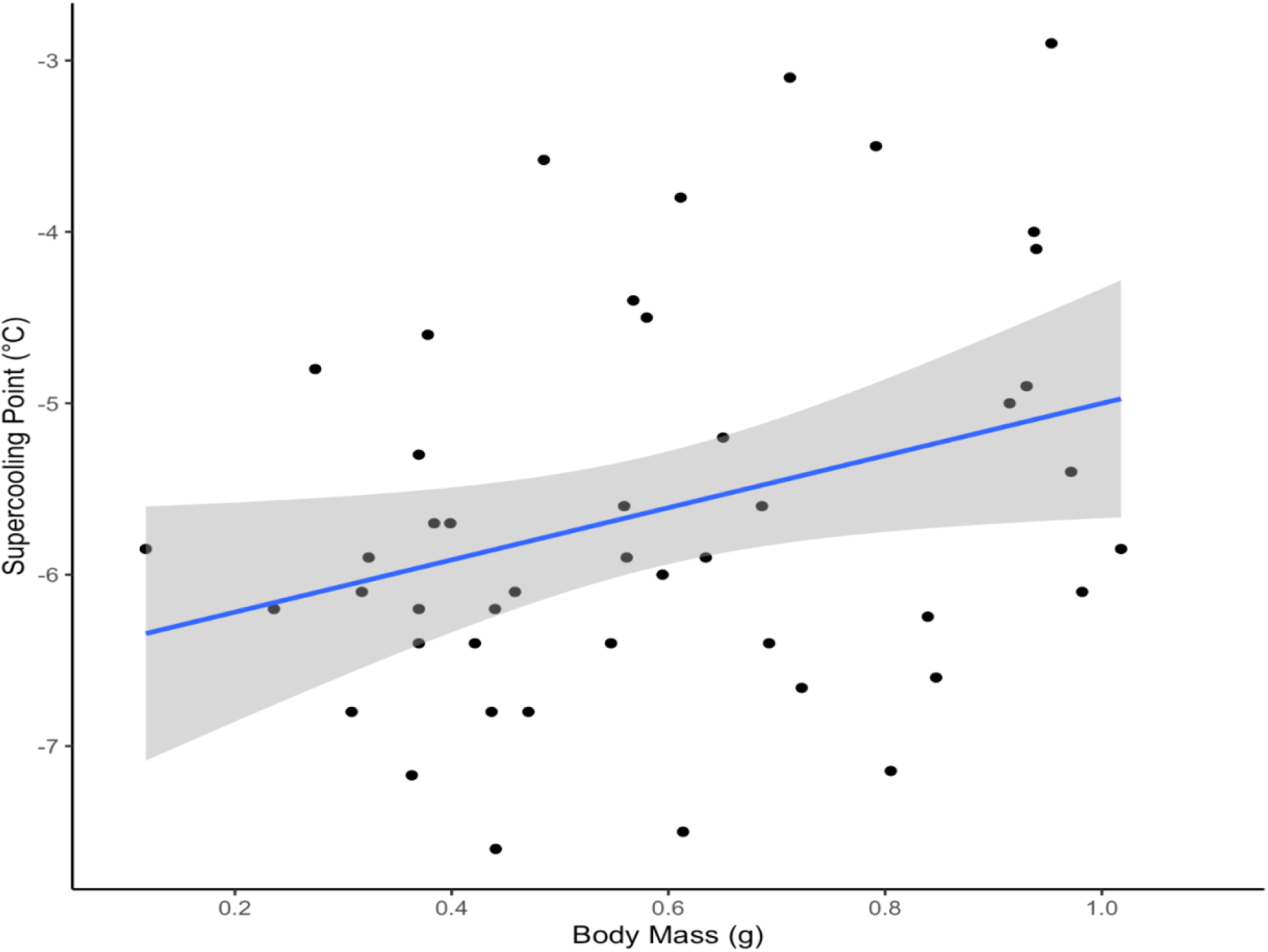
Slug supercooling point increases with increasing wet body mass. Linear trendline is dark blue with grey shaded standard error. N = 48, p = 0.0224.

### Body water content

Wet mass of *A. valentianus* was measured right after they were taken from their acclimation chambers, and dry mass was measured after slugs had been freeze dried. Their body water content is expressed as a percentage (1 - (dry mass / wet mass) x 100), a higher percentage meaning the slug had a higher body water content at the end point of acclimation. Slugs acclimated to 15° C temperatures had significantly lower body water content (83.68 ± 0.0040%) than those acclimated to 20 °C (85.52 ± 0.0038%; df = 1, 49, p = 0.0027). Body water content was not affected by photoperiod (df = 1, 49, p = 0.41). The interaction term showed the effect of acclimation photoperiod on body water content was affected by acclimation temperature, whereby slugs at high temperature and short days had lower body water content (df = 1, 49, p = 0. 044).

### NMR metabolomics

Proton NMR analysis identified 47 metabolites in *A. valentianus*. Of these 47, four were alcohols and polyols, 11 were organic acids, 25 were amino acids, peptides, and analogues, two were sugars, and five were other metabolites (Table 1). After controlling for false discovery rate (FDR), the concentrations of three metabolites (formate, L-glutamine, and L-threonine) were significantly affected by slug acclimation conditions. A two-way ANOVA showed that formate concentrations were affected only by temperature and were depressed at short day 15 °C acclimation conditions relative to short day 20 °C (p = 0.0045, df = 1, 32; Fig. 5). Formate concentrations were not significantly affected by day length, and the interaction term between acclimation temperature and day length was also not significant (day length: p = 0.65, df = 1, 32; day length × temperature: p = 0.44, df = 1, 32). L-glutamine concentrations were affected by temperature and photoperiod and were depressed at short day 15 °C acclimation conditions relative to short day 20 °C and long day 20 °. (temperature: p = 0.00056, df = 1, 32; day length: p = 0.17, df = 1, 32; Fig. 5). The interaction term between acclimation temperature and day length for L-glutamine was not statistically significant (day length × temperature: p = 0.31, df = 1, 32). L-threonine concentrations were significantly affected by acclimation day length (p = 0.0015, df = 1, 32, Fig. 5). The interaction term also showed that the effect of daylength was more pronounced under certain temperatures, whereby slugs acclimated to long day conditions at 15 °C significantly increased L-threonine concentrations relative to the other acclimation conditions, but that acclimation temperature by itself did not significantly effect concentrations (day length × temperature: p = 0.0037, df = 1, 32; temperature: p = 0.40, df = 1, 32; Fig. 5).

**Fig. 5.**
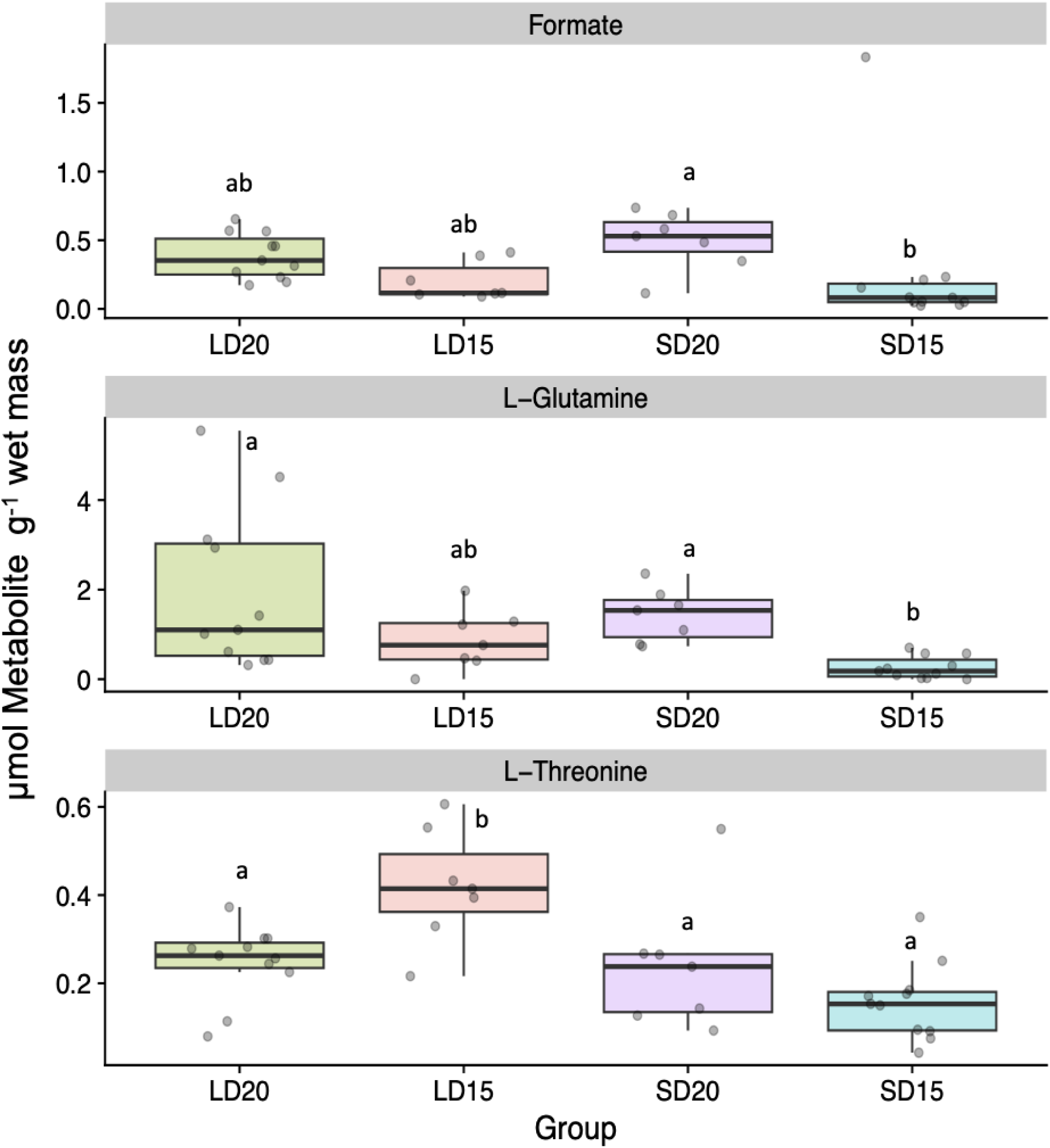
Formate, L-Glutamine, and L-Threonine amounts in *A. valentianus* after three weeks of acclimation at one of four conditions: long day (LD) at 20 °C, LD at 15 °C, SD at 15°C, or short day (SD) at 20 °C. Amounts are normalized by individual slug wet mass. The bottom and top of the box corresponds to the first and third quartiles, the median is denoted by the line within the box, and the whiskers extend to the largest or smallest value in the group (or no further than 1.5 * interquartile region). The underlying grey points represent the values of an individual sample. n = 7-11, letters denote significance between groups (statistics were completed on square root transformed and auto-scaled data).

**Table 1.** List of metabolites identified in *A. valentianus*. The mean and standard error of the total amounts (µmol) found for each metabolite is indicated in brackets [mean±SE].

| Amino acids | Organic Acids | Other metabolites | Peptides and Analogues | Sugars, alcohols and polyols |
| --- | --- | --- | --- | --- |
| 1-Methylhistidine [0.25±0.04] | 2-Hydroxybutyric acid [0.36±0.03] | Acetone [0.02±0] | Aspartate [0.71±0.14] | D-Glucose [9.43±1.07] |
| Glycine [0.52±0.1] | 3-Hydroxybutyric acid [0.14±0.01] | Carnitine [0.13±0.03] | Betaine [2.11±0.16] | Ethanol [0.15±0.02] |
| Isoleucine [0.18±0.02] | Acetic acid [0.1±0.01] | Urea [7.42±1.25] | Choline [0.09±0.01] | Glycerol [2.78±0.39] |
| L-Alanine [0.09±0.01] | Acetoacetate [0.05±0.01] |  | Creatine [0.13±0.02] | Isopropyl alcohol [0.03±0] |
| L-Arginine [0.79±0.08] | Citric acid [0.12±0.02] |  | Creatinine [0.08±0.01] | Methanol [0.92±0.19] |
| L-Asparagine [0.6±0.17] |  |  | Dimethyl sulfone [0.11±0.03] |  |
| L-Glutamic acid [1.98±0.16] |  |  | Formate [0.33±0.06] |  |
| L-Glutamine [1.12±0.21] |  |  | Isobutyric acid [0.01±0] |  |
| L-Histidine [0.23±0.03] |  |  | L-Lactic acid [0.21±0.11] |  |
| L-Leucine [0.42±0.04] |  |  | Malonate [0.39±0.11] |  |
| L-Lysine [0.2±0.03] |  |  | Methionine [0.14±0.02] |  |
| L-Ornithine [0.2±0.03] |  |  | Pyruvic acid [0.02±0] |  |
| L-Phenylalanine [0.04±0] |  |  |  |  |
| L-Proline [0.87±0.1] |  |  |  |  |
| L-Serine [1.85±0.18] |  |  |  |  |
| L-Threonine [0.25±0.02] |  |  |  |  |
| Succinate [0.07±0.01] |  |  |  |  |
| Tryptophan [0.05±0.01] |  |  |  |  |
| Valine [0.1±0.01] |  |  |  |  |

Long day (LD) 20 °C and short day (SD) 15 °C acclimated slugs were chosen as samples to compare with pathway analysis in MetaboAnalyst 6.0, as they were the most extreme groups. Four metabolic pathways were significantly enriched between long day (LD) 20 °C acclimated slugs and short day (SD) at 15 °C acclimated slugs (Table 2 and Fig. 6): glyoxylate and dicarboxylate metabolism, purine metabolism, glycine, serine and threonine metabolism (Fig. S4), and cysteine and methionine metabolism.

**Fig. 6.**
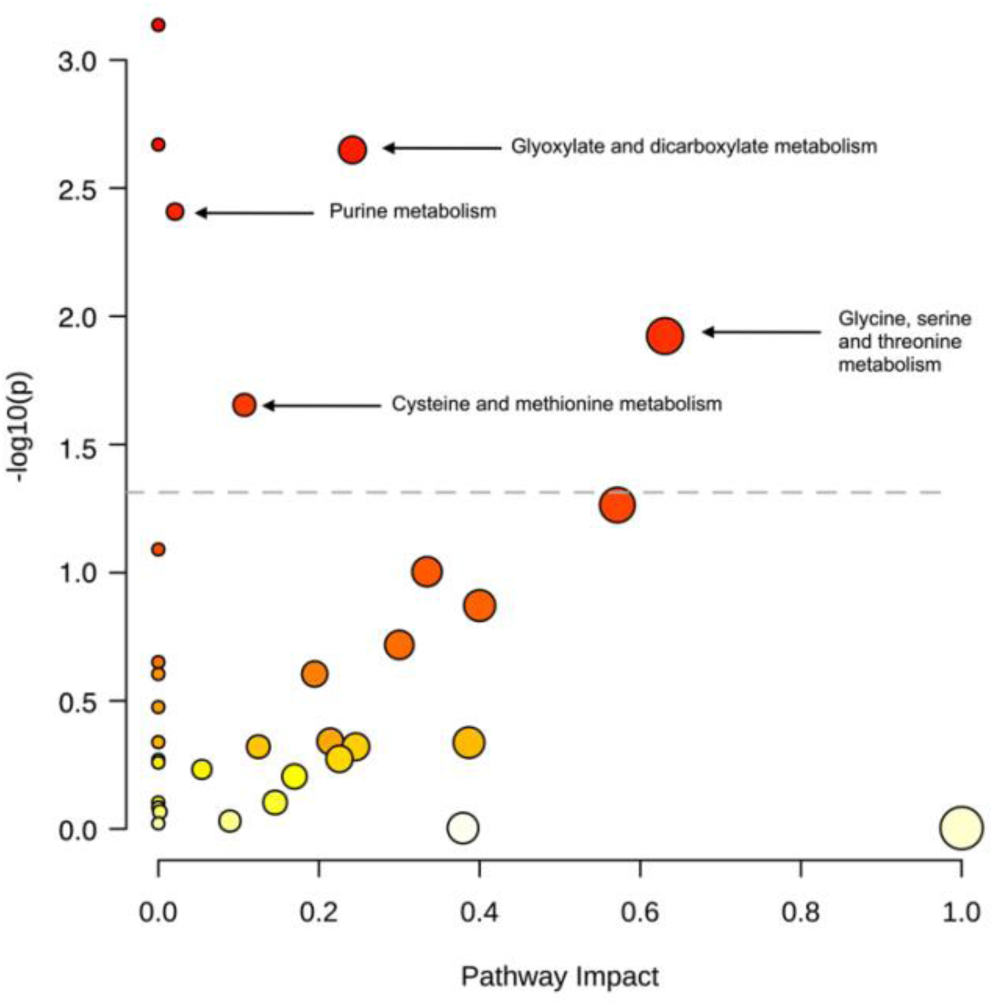
The results of pathway analysis of the metabolomic data based on *Drosophila melanogaster* Kyoto Encyclopedia of Genes and Genomes (KEGG) pathways. Larger circles represent greater pathway enrichment, and darker circle colours (yellow being lighter, red being darker) represent lower p-values. Dashed grey line indicates p = 0.05. Pathways that have an impact factor >0.0 and a p<0.05 are labelled with arrows.

**Table 2.** Pathway analysis of metabolic change in *A. valentianus* following acclimation to SD15 conditions relative to LD20 conditions. The analyses were conducted by the pathway analysis modules of MetaboAnalyst 6.0. Metabolite concentrations were normalized by slug wet weight, square root transformed and then auto-scaled. Pathways that have an impact factor >0.0 and a p<0.05 are included below. ‘Match Status’ refers to the number of metabolites altered in the pathway, ‘p’ is the p-value, ‘FDR’ is the false discovery rate, and ‘Impact’ denotes the impact factor which is a MetaboAnalyst derived value calculated by adding up the importance measures of each of the matched metabolites and then dividing by the sum of the importance measures of all metabolites in each pathway.

| Pathway Name | Match Status | p | -log(p) | Holm p | FDR | Impact |
| --- | --- | --- | --- | --- | --- | --- |
| Glyoxylate and dicarboxylate metabolism | 6/24 | 0.00224 | 2.649 | 0.0718 | 0.0196 | 0.242 |
| Purine metabolism | 2/68 | 0.00391 | 2.408 | 0.121 | 0.0273 | 0.0208 |
| Glycine, serine and threonine metabolism | 5/29 | 0.0119 | 1.923 | 0.358 | 0.0696 | 0.631 |
| Cysteine and methionine metabolism | 3/32 | 0.0222 | 1.654 | 0.643 | 0.1108 | 0.107 |

## Discussion

Here we show that *A. valentianus* collected from Kyoto, Japan survives cold better when acclimated to shorter photoperiods. However, this species does not exhibit significant freeze tolerance. We also found evidence that some low molecular weight molecules (L-glutamine, L-threonine and formate) were significantly affected by varying photoperiod and temperature acclimation, but there was no evidence of polyol or sugar accumulation.

Therefore, the slug *A. valentianus* displays plastic cold tolerance without accumulating low molecular weight cryoprotectants like some other invertebrate species, such as insects. Overall, investigating the temperature tolerances and environmental factors that allow invasive molluscs to establish can help deepen our understanding of invasive species biology.

### Environmental factors

We aimed to determine how environmental factors, specifically the two main cues that prepare terrestrial molluscs for winter 1) shortening day lengths and 2) cooling temperatures effect the acquisition of cold and freeze tolerance. Given that these cues indicate future winter conditions, we anticipated that slugs exposed to shorter photoperiods and lower temperatures may physiologically prepare for winter, and thus be more resilient to sudden onsets of sub-zero temperatures. We found that when *A. valentianus* were acclimated to short day conditions (12L:12D), they were able to survive a short, partial freeze exposure significantly better compared to when they had been acclimated to long day conditions (16L:8D; Fig. 1). We did not see the same effect with low temperature acclimation, which suggests that shortening day lengths may be the driving factor behind seasonal acquisition of cold tolerance.

While this is the first study to examine specifically the freeze tolerance of *A. valentianus* under different acclimation conditions, there have been previous studies examining the seasonal plasticity of cold tolerance of *A. valentianus* and other terrestrial slug species. The enhancement of cold tolerance in *A. valentianus* begins in the fall before the coldest winter temperatures set in (Udaka et al., 2007). *Ambigolimax valentianus* collected in fall and winter from October – February in Kyoto, Japan were more cold tolerant than those collected throughout the spring and summer months (Udaka et al., 2008). Lab-raised *A. valentianus* also exhibited increased cold tolerance with both short day conditions and lower acclimation temperatures (Udaka et al., 2008). Taken together with our results, this may suggest that developmental acclimation to temperature and photoperiod may be important for this species, and that photoperiod may take precedence to influence seasonality in their adult stage.

The evidence that low temperature acclimation can induce increased cold tolerance in terrestrial molluscs is mixed. For example, two other terrestrial species of slug (*D. reticulum* and *Arion hortensis*) did not show any change in cold tolerance after short term acclimation to 5 °C (Mellanby, 1961). The land snail *Helix aspera* also did not change its freeze tolerance in response to lower temperature acclimatization, but showed an increase in freeze tolerance due to shortening photoperiods (Ansart et al., 2001). However, several other species of land snails (*D. cronkhitei, A. alternata,* and *G. armifera*) have exhibited increased freeze tolerance (decreased SCPs) after exposure to low acclimation temperatures (Riddle & Miller, 1988). Taken together, this indicates that photoperiod may be a stronger cue than low temperatures for some species of terrestrial molluscs to induce seasonal cold or freeze tolerance, including our study species *A. valentianus*.

### Freeze tolerance

Based on the evidence from this study, we conclude that *A. valentianus* is not strongly freeze tolerant, however, determining it as such requires several important considerations. The survival of *A. valentianus* following brief periods of freezing suggests some level of resilience to ice formation, as observed in Fig. 1 where short-day-acclimated slugs exhibited approximately 40% survival after spending seconds being frozen. Whether an individual survived this short freeze exposure may have been dependent on where ice initially formed (ice forming in the head could be more damaging than ice growth in the posterior end of the body). Slugs could survive exposure to cold temperatures without freezing, but mortality depended on whether freezing actually occurred (Fig. 2). This indicates that low temperatures are likely not the cause of death, but rather the presence and extent of ice within the body. It is important to highlight that the observed differences in freezing survival between the two experiments can be attributed, at least in part, to variations in the freezing protocols. Specifically, the slugs used in Fig. 2 spent an average of 35 minutes in the freezing bath, with varying amounts of time spent frozen. In contrast, the slugs in Fig. 1 spent an average of 20 minutes in the freezing bath and were only frozen for a few seconds. Taken together, these two experiments indicate that *A. valentianus* can survive an extremely limited amount of ice formation (probably dependent on total ice content and ice forming in a location where they can tolerate it) but would likely not be able to experience a biologically relevant freeze exposure, and thus are likely not freeze tolerant.

Beyond direct freezing experiments, other physiological factors can inform the classification of an organism as freeze-tolerant, such as changes in SCP, changes in body water content, and differences in body size. Freeze tolerant animals often demonstrate increased SCPs as ice nucleators are accumulated (Toxopeus & Sinclair, 2018). We found that *A. valentianus* SCPs did not change with temperature or photoperiod acclimation in a statistically significant way, even though slugs acclimated to short day 15 °C conditions had slightly higher SCPs than long day 20 °C slugs (difference = 0.90 °C; Fig. 3). However, because this difference was less than 1 °C, and the error range on the thermocouples themselves is about ± 1 °C, this difference is likely not biologically significant. Freeze tolerant invertebrates will often increase their SCP during the winter (Kennedy et al., 2020). Because there was no significant increase in the SCP of *A. valentianus*, they may not be accumulating ice nucleators as winter approaches, lending support to the idea that this species may not be meaningfully freeze tolerant. Additionally, we found that SCP varied with slug weight, with heavier slugs having a higher SCP compared to lighter slugs (Fig. 4). This is a well-understood phenomenon as heavier slugs have more water molecules, and thus a higher probability that any one of those molecules freeze during a cold exposure (Bigg, 1953; Zachariassen & Kristiansen, 2000). This likely means that smaller *A. valentianus* individuals may have a better chance of surviving sub-zero temperatures during the winter, especially since freezing seems to almost instantly induce mortality. On a broader level, *A. valentianus* is generally a smaller species of slug (mean ± SE = 0.59 ± 0.04 g, range = 0.12–1.26 g;). Body size influences how terrestrial molluscs respond to freezing conditions - smaller species tend to avoid freezing, while larger ones are more likely to tolerate freezing (Ansart & Vernon, 2003). Finally, we found that *A. valentianus* decreased their body water content after acclimation to 15° C. Freeze tolerant animals are known to decrease their body water in preparation for freezing which reduces the risk of intracellular ice formation and limits mechanical damage within the body due to excess ice formation (Storey & Storey, 2013). However, freeze avoidant animals do this too. Although we saw evidence of a body water decrease, it was only very slight (85% body water in 20 °C conditions vs 83.5% body water in 15 °C conditions), and probably not biologically significant. Overall, *A. valentianus* did not display much difference in SCP or body water content between acclimation conditions and is a relatively smaller species, which supports the conclusion that they are likely not freeze tolerant.

While we did not find evidence of freeze tolerance in our study, it is possible that this conclusion is limited by our sampling. First, there are populations of *A. valentianus* that live in places that experience sub-zero temperatures often throughout the winter like Sweden, Finland, western Canada, southern Argentina and northern Japan (*Ambigolimax valentianus,* GBIF). However, the population of *A. valentianus* individuals in this experiment were collected from Kyoto, Japan where it rarely gets below freezing (the average historical minimum temperatures in nearby Osaka only reach 3 °C in the December, the coldest month of the year; Japan Meteorological Agency, 2024). Furthermore, these slugs were collected in June and July in Kyoto, where the historical maximum temperatures reach 31.8 °C and 33.7 °C in each month respectively (Japan Meteorological Agency, 2024). Despite not experiencing sub-zero temperatures naturally and being collected in the summer, these slugs were still able to survive over 30 minutes of exposure to -6 °C following a three-week acclimation to 15 °C, as long as the body fluids did not freeze (Fig. 2). Remarkably, winter-collected *A. valentianus* show an even greater thermal tolerance, with a lower lethal temperature of -8 °C in January (Udaka et al., 2008). Although this population of *A. valentianus* was collected from a warm region during a warm time of year and may not represent the most cold-adapted population of the species, it still demonstrated the ability to tolerate subsequent exposure to relatively low temperatures.

In general, invasive species often have a broad thermal width, and therefore have the ability to function physiologically across a wide range of temperatures (Kelley, 2014). Here we see that *A. valentianus* is cold tolerant, but furthermore that their plasticity to develop cold tolerance after acclimation is higher than one may expect from their environment. These traits might be the key to its invasive success. Additionally, low temperatures are a key factor that prevent species from invading a new region (Jarošík et al., 2015; Stachowicz et al., 2002). This slug species can tolerate cold temperatures and therefore may be less constrained by low temperature limits, meaning this species could potentially establish itself in a broader range of environments. For example, the invasive Asian lady beetle *Harmonia axyridis* has a lower critical thermal minimum than its native South American counterpart, enabling it to expand further south into colder regions of Chile (Boher et al., 2018). This may explain why *A. valentianus* is not only invasive in Kyoto, Japan, but other areas like Scandinavia where winters are much colder.

Intraspecific variation in thermal tolerance has been well documented in several species of insects including fruit flies and ants (Sarup et al., 2009; Sgrò et al., 2010; Tonione et al., 2020). For intertidal mollusc species, thermal tolerances can vary even on the scale of shore heights, with animals higher on the shore often tolerating more extreme temperatures (Kennedy et al., 2020; Sokolova et al., 2000). In terrestrial molluscs, two species of land snails, *Helix pomatia* and *Cornu aspersum*, show intraspecific variation in cold tolerance, with those from colder climatic conditions exhibiting an enhanced supercooling capacity (Ansart & Vernon, 2004; Nicolai et al., 2005). However, it remains uncertain if this variation was driven by climatic conditions or differences in body size and water content between populations. *Ambigolimax valentianus* may also display intraspecific thermal tolerance, where populations that live in places with colder winters displaying enhanced cold and/or freeze tolerance. Although purely speculative, it is possible that this population of *A. valentianus* residing in a warmer region of Japan, does not exhibit freeze tolerance, whereas a population living in Sweden would, however there are no investigations that we know of confirming freeze tolerance in *A. valentianus* from cold areas. To definitively determine whether *A. valentianus* is freeze-tolerant, future studies could replicate this experiment using a population that regularly encounters sub-zero conditions during winter.

Although *A. valentianus* is cold tolerant, it still faces mortality if it freezes. As a result, these slugs must adopt alternative strategies to cope with winter conditions. Field observations have noted that *A. valentianus* are less reproductively active during the coldest months of the year (Udaka et al., 2007), indicating possible overall reductions in activity. We also observed huddling behaviour in *A. valentianus* in response to winter conditions (**Error! Reference source not found.** S5), which is a behavioural response to reduce evaporative water loss (A. C ook, 1981). When faced with freezing temperatures *A. valentianus* may also behaviourally thermoregulate by moving to a place where they are not in contact with the frozen ground, thus avoiding freezing all together.

### Metabolites

If shortening day lengths and cooling temperatures signal that winter is approaching and the risk of freezing is imminent, then we may expect that *A. valentianus* metabolites may alter in response to oncoming cold temperatures. In freeze tolerant species, specific metabolites may function as low molecular weight cryoprotectants by balancing osmotic ratios during freezing and other cryoprotective functions. For example, insects are known accumulate multimolar quantities of metabolites such as sugars, polyols, and amino acids to act as cryoprotectants in cold conditions (Lee, 2010; Toxopeus & Sinclair, 2018). Accumulating metabolites increases the osmolarity of body fluids, thereby depressing the organism’s supercooling point, prevent intracellular freezing, and protect from cellular dehydration (Storey, 1997). These metabolites may function on a colligative basis, by increasing the metabolite pool concentration to enhance freeze tolerance, or on a non-colligative basis, where each metabolite has a unique protective function (Toxopeus & Sinclair, 2018).

Our NMR analysis identified 47 metabolites in *A. valentianus* including organic acids, sugars, and amino acids, among others (Table 1). Among these, three were changed with acclimation conditions: L-glutamine, L-threonine and formate. Notably, glucose concentrations were not altered with shortening photoperiods and lower temperatures. This is unlike freeze tolerant species such as vertebrates, annelids, and other phyla that can upregulate glucose on the molar level in response to sub-zero temperatures (Storey, 1997). Here, we show that *A. valentianus* has micromolar level quantities of glucose (on average 9.43 µmol per animal; Table 2). This baseline amount of glucose is consistent with what has been observed in other terrestrial slug species, such as *A. rufus*, *D. leave*, and *Arion lusitanicus*, which also maintain glucose in the micromolar range (Slotsbo et al., 2012; Storey et al., 2007).

However, there is a key difference: unlike *A. valentianus*, these other slug species significantly increase their glucose concentrations when exposed to cold or freezing conditions (Slotsbo et al., 2012; Storey et al., 2007). We found that glucose concentrations in *A. valentianus* did not increase in response to winter conditions. Other gastropods, such as the land snail *Helix pomatia* also show no such accumulation of glucose in response to winter conditions such as shortening photoperiods or cooling temperatures (Nowakowska et al., 2011). Even though glucose did not differ between acclimation conditions in *A. valentianus*, there were other metabolites that were significantly altered.

We saw decreased concentrations of L-glutamine and formate in slugs acclimated to short day 15 °C conditions (Fig. 5). Glutamine is an amino acid that has many important functions in the body. Glutamine can be synthesized into glutamate in one reaction, and both glutamine and glutamate are metabolic intermediates (Labow & Souba, 2000). Glutamine concentrations have also been shown to decrease strongly during anoxia and freezing in two other species of terrestrial slug, *D. reticulatum* and *D. leave* (Storey et al., 2007). However, some cold acclimated insects show increases in glutamine concentrations such as wheat weevils (*Sitophilus granaries*; Fields et al., 1998). Formate is the conjugate base of formic acid that interacts with energy metabolism. Both of these metabolites were decreased in cold acclimated *A. valentianus,* suggesting that there may be metabolic suppression during the winter.

We also found that L-threonine was upregulated under long day photoperiods and 15 °C conditions (Fig. 5). Threonine is an amino acid that plays and important role for cell growth, protein synthesis, and lipid metabolism (Tang et al., 2021). Threonine concentrations increase in wheat midge larvae before diapause, and then decrease during diapause (Huang et al., 2022). Perhaps this intermediate acclimation condition caused *A. valentianus* to upregulate threonine. *Ambigolimax valentianus* mucus contains a high percentage of glycine, which is downstream from its precursor, threonine (Li & Graham, 2007). We found that glycine, serine, and threonine metabolism was altered by acclimation condition (Fig. S4). When approaching fall conditions, *A. valentianus* may alter its mucus production which could be reflected in its metabolite concentrations.

Our findings show that metabolites like L-glutamine and formate are strongly reduced in response to winter seasonal cues, suggesting possible whole body metabolic suppression leading up to winter. It is unlikely that metabolites act as low molecular weight cryoprotectants in *A. valentianus* and are likely not key contributors to cold or freeze tolerance in this organism. *Ambigolimax valentianus* exhibited some minimal freeze tolerance in short day 20 °C conditions, even though there were no differences in metabolite concentrations compared to our summer acclimated slugs, which did not display any freeze tolerance and limited cold tolerance. We also did not detect any differences in glucose, a typical cryoprotectant that has been observed in other slug species in response to freezing (Slotsbo et al., 2012). In general, we did not observe many large variations in metabolite concentrations within our acclimation groups, with only 3 out of 47 detected metabolites showing any differences. Therefore, other mechanisms likely drive the cold tolerance of *A. valentianus*.

If small molecules are not contributing significantly to the acquisition of cold tolerance in *A. valentianus*, then there could be other mechanisms of cold tolerance at play. The heat shock proteins (HSPs) are a diverse group of chaperone proteins that are produced in response to many types of physiological stressors including heavy metal contamination, disease, osmotic stress, heat shock, and cold stress (Encomio and Chu, 2005; Franzellitti and Fabbri, 2005; Singer et al., 2005; Snyder et al., 2001). Although not classically considered cryoprotectants, HSPs are a crucial part of freeze tolerance in yeast, fruit flies and the beet armyworm (Choi et al., 2014; King and Macrae, 2015; Pacheco et al., 2009; Xue et al., 2009). This may be especially important for the invasive *A. valentianus*, since non-native species often have greater expression of HSPs, which enhances their temperature tolerance (Kelley, 2014).

Another way in which organisms survive the cold is through homeoviscous adaptation. As temperatures decrease, the fluidity of cellular membranes declines, leading to potential physiological disruptions due to reduced membrane protein activity and energy availability (de Mendoza and Cronan, 1983; Somero et al., 2017). To counteract this, many organisms undergo homeoviscous adaptation, altering their lipid composition to preserve membrane fluidity in cold conditions (Sinensky, 1974). This adaptation has been studied in various species including *Drosophila melanogaster* (MacMillan et al., 2009), and could play a role in the cold tolerance of *A. valentianus*.

Finally, ice binding proteins (IBPs) may play a role in *A. valentianus* freeze tolerance. IBPs are used by freeze-avoidant species to prevent ice formation through a process called thermal hysteresis, and this process has been observed in freeze-avoidant intertidal and terrestrial species (Hargens & Shabica, 1973; Hawes et al., 2010; Tyshenko et al., 1997; Zachariassen & Kristiansen, 2000). Although metabolites may not significantly contribute to the cold tolerance of *A. valentianus*, it is possible that other mechanisms like HSPs, lipid membrane modifications, or IBPs are involved, warranting further investigation in future studies.

## Conclusions

Our study identified that *A. valentianus* is likely not freeze tolerant but does exhibit some plasticity in cold tolerance in response to shortening photoperiods and low temperature acclimation. This species is invasive and is considered a pest in residential gardens and agricultural settings around the world. The reasons for the mass occurrences of *A. valentianus* are not fully untangled, though its success may be due to its impressive thermal tolerance to hot summers and low winter temperatures, potentially allowing it to outcompete other native slug species. This study in particular aims to understand a neglected aspect of molluscan physiology, low temperature tolerance, to better understand the dispersal patterns and potential invasive range of *A. valentianus*. Low temperature tolerances are of particular importance as climate warming is resulting in poleward range shifts meaning that cold and/or freeze tolerance is an important limiting factor at poleward range edges of both native and non-native species (Jarošík et al., 2015). If invasive species have a superior ability to tolerate freezing and can outcompete native species in the face of climate change, our ecosystems may be permanently altered. Therefore, understanding cold tolerance of species, both native and invasive, is an important part of understanding potential range spread.

## Acknowledgements

The authors would like to thank Chris Harley and Patricia Schulte for their thoughtful feedback on the manuscript. They would also like to thank the UBC NMR facility with assistance processing the NMR samples.

## Data Availability

Data and code is available in a public repository: https://github.com/laurentjgill/slug

