## Supplemental figures for "The cold tolerance of the terrestrial slug, *Ambigolimax valentianus*"

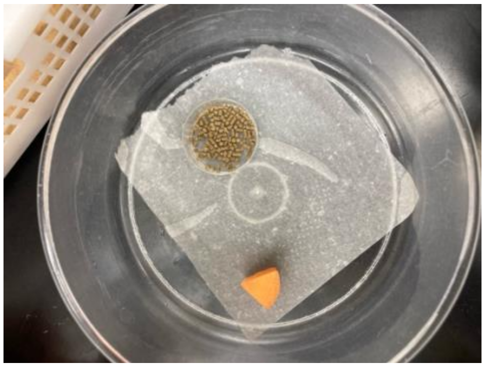

Supplemental Fig. 1: Plastic containers where slugs were kept.

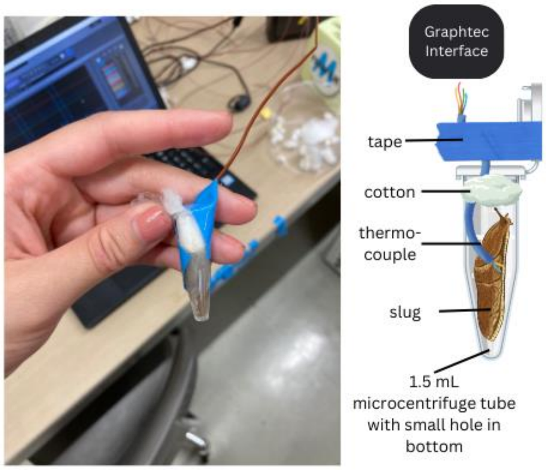


Supplemental Fig. 2 Configuration of slugs and thermocouples in microcentrifuge tube before freeze exposure. Graphics sourced from Canva.

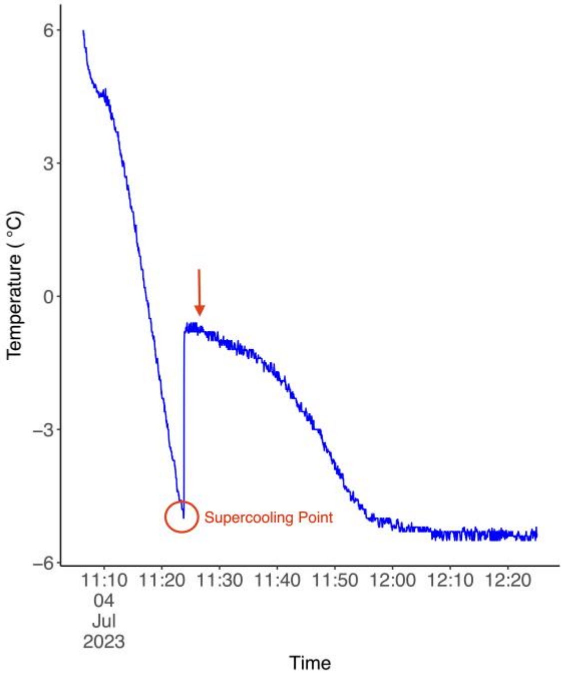

Supplemental Fig. 3 An example of a slug's temperature trace during an exposure to low temperature. Slugs were placed in a refrigerated circulating bath at a starting temperature of 5 °C, then frozen with a cooling rate of -0.75 °C per minute, until the bath reached -7 °C. The red circle refers to the supercooling point - the lowest temperature reached before the exothermic release of energy caused by ice formation. The red arrow refers to the point where slugs were removed in the survival experiment.

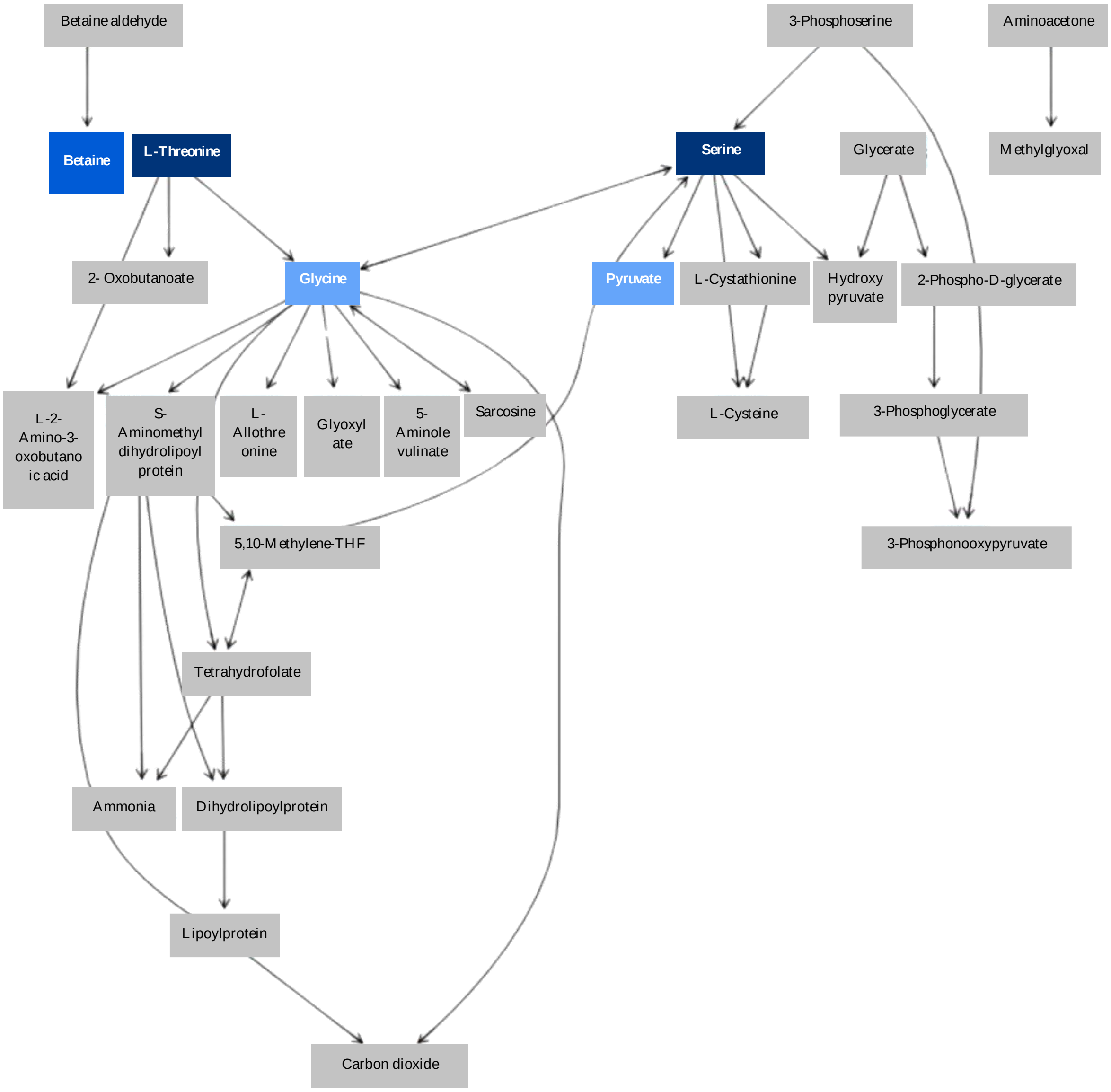


Supplemental Fig. 4 Glycine, serine and threonine metabolism. Grey boxes represent metabolites that were not measured through NMR analysis, and coloured boxes represent measured metabolites. Darker coloured boxes indicate the metabolite was more significantly altered between SD15 and LD20 conditions. Adapted from MetaboAnalyst.

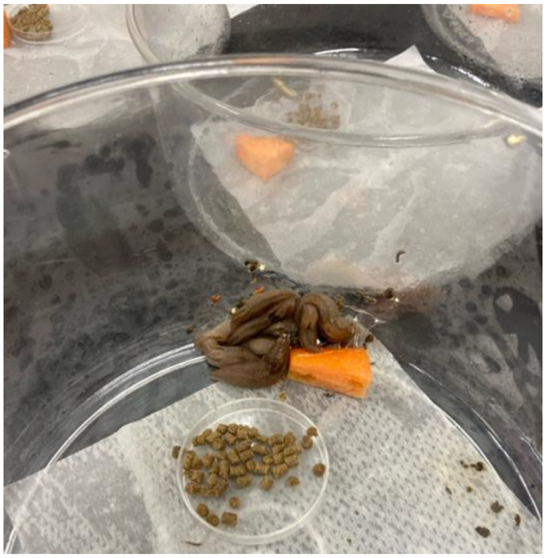

Supplemental Fig. 5 A group of *A. valentianus* individuals displaying huddling behaviour.
